# Actomyosin forces and the energetics of red blood cell invasion by the malaria parasite *Plasmodium falciparum*

**DOI:** 10.1101/2020.06.25.171900

**Authors:** Thomas C. A. Blake, Silvia Haase, Jake Baum

## Abstract

All symptoms of malaria disease are associated with the asexual blood stages of development, involving cycles of red blood cell (RBC) invasion and egress by the *Plasmodium* spp. merozoite. Merozoite invasion is rapid and is actively powered by a parasite actomyosin motor. The current accepted model for actomyosin force generation envisages arrays of parasite myosins, pushing against short actin filaments connected to the external milieu that drive the merozoite forwards into the RBC. In *Plasmodium falciparum*, the most virulent human malaria species, Myosin A (PfMyoA) is critical for parasite replication. However, the precise function of PfMyoA in invasion, its regulation, the role of other myosins and overall energetics of invasion remain unclear. Here, we developed a conditional mutagenesis strategy combined with live video microscopy to probe PfMyoA function and that of the auxiliary motor PfMyoB in invasion. By imaging conditional mutants with increasing defects in force production, based on disruption to a key PfMyoA phospho-regulation site, the absence of the PfMyoA essential light chain, or complete motor absence, we define three distinct stages of incomplete RBC invasion. These three defects reveal three energetic barriers to successful entry: RBC deformation (pre-entry), mid-invasion initiation, and completion of internalisation, each requiring an active parasite motor. In defining distinct energetic barriers to invasion, these data illuminate the mechanical challenges faced in this remarkable process of protozoan parasitism, highlighting distinct myosin functions and identifying potential targets for preventing malaria pathogenesis.

## Introduction

Malaria disease is caused by single-celled, obligate intracellular parasites from the genus *Plasmodium*, the most virulent species being *Plasmodium falciparum*. All symptoms of malaria disease result from cycles of parasite invasion into, development within and egress from red blood cells (RBCs) so improved understanding of the process of parasite invasion remains a central target for future therapeutics (Burns *et al*, 2019). RBC invasion is mediated by merozoites, specialised motile cells around 1 μm in size (Dasgupta *et al*, 2014) that employ substrate-dependent gliding motility (Yahata *et al*, 2020). Merozoites are primed for invasion by phosphorylation of the motility apparatus before RBC egress (Alam *et al*, 2015) and have a short window of viability to invade, in the seconds to minutes range (Boyle *et al*, 2010). Having encountered an RBC, merozoites initially attach to the RBC membrane via weak, non-specific interactions followed by strong contact via parasite adhesins (Tham *et al*, 2015). Once attached, the process of invasion is rapid. Video microscopy of invasion reveals that it takes 20-30 s (Dvorak *et al*, 1975; Gilson & Crabb, 2009) and consists of several distinct phases. The merozoite actively deforms the RBC, reorientates to its apex, and then attaches irreversibly to the RBC via formation of a tight junction (TJ) (Riglar *et al*, 2011), a parasite secreted complex thought to act as a point of traction (Baum & Cowman, 2011). Active penetration of the RBC then follows (Miller *et al*, 1979). Parasite adhesins are secreted from apical organelles, the micronemes and rhoptries, in response to a signalling cascade involving Ca^2+^ ions (Singh *et al*, 2010; Bullen *et al*, 2016) which also regulates phosphorylation of adhesins and actomyosin components (Paul *et al*, 2015; Fang *et al*, 2018). Finally, after completion of invasion the RBC usually undergoes a process of echinocytosis, shrinking and producing spicules in response to the perturbation to osmotic balance that follows rhoptry secretion and membrane disruption.

*Plasmodium* merozoites rely on a conserved molecular motor for gliding and invasion, centred around a MyoA motor complex (MMC) (Baum *et al*, 2006) or glideosome, situated in the narrow space between the parasite plasma membrane and the double membrane inner membrane complex (IMC) (Frénal *et al*, 2017). *P. falciparum* MyoA (PfMyoA) is a small, atypical myosin motor that belongs to the alveolate-specific class XIV. Like other class XIV myosins, PfMyoA comprises a motor domain and a light chain-binding neck domain but lacks an extended myosin tail domain for cargo-binding. Instead, PfMyoA relies on a regulatory light chain or myosin tail interacting protein, MTIP, that binds the extreme end of the neck domain and unusually possesses a long, disordered N-terminal domain to anchor PfMyoA to the IMC (Bergman *et al*, 2003). Maximal *in vitro* activity of PfMyoA requires the binding of MTIP and a second essential light chain, PfELC (Bookwalter *et al*, 2017), recently shown to stabilise the PfMyoA neck domain and is critical for parasite replication (Moussaoui *et al, manuscript submitted*). The complex of PfMyoA and its light chains (the PfMyoA triple complex) is anchored to the IMC by a glideosome associated protein, PfGAP45 (Frénal *et al*, 2010; Perrin *et al*, 2018) and other GAP proteins embedded in the IMC membranes that together complete the MMC. Genetic demonstration that *Plasmodium berghei* MyoA is critical for motility of the mosquito-infecting ookinete stage (Sidén-Kiamos *et al*, 2011) and PfMyoA is critical for blood-stage replication (Robert-Paganin *et al*, 2019) confirm that PfMyoA is at the core of the parasite force generation and hence invasion/motility machinery.

PfMyoA produces directional force by undergoing cycles of ATP hydrolysis and conformational change, allowing it to translocate short, unstable actin filaments (Das *et al*, 2017; Lu *et al*, 2019) that are in turn connected to the external substrate. The crystal structures of the PfMyoA motor domain (Robert-Paganin *et al*, 2019) and triple complex (Moussaoui *et al, manuscript submitted*) reveal a unique mechanism of force production involving stabilisation of the rigor-like state by an interaction between K764 in the converter and phospho-S19 in the N-terminal extension (NTE). Disruption of this interaction *in vitro* reduced the velocity of PfMyoA but increased its maximal force production (Robert-Paganin *et al*, 2019), suggesting that phosphorylation of S19 is required for maximal myosin velocity. A “phospho-tuning” model was therefore proposed to explain how the same motor is optimised for speed in fast gliding stages or force production during invasion.

Several questions remain about PfMyoA organisation and function, in particular how motor force is integrated with retrograde flow of parasite plasma membrane (Quadt *et al*, 2016; Moreau *et al*, 2017; Whitelaw *et al*, 2017; Gras *et al*, 2019) and is applied across the parasite (Tardieux & Baum, 2016). Evidence from imaging suggests MyoA may regulate force production at discrete adhesion sites, rather than acting as a simple linear motor (Münter *et al*, 2009; Whitelaw *et al*, 2017). However, these questions have been addressed in non-merozoite stages of *Plasmodium* or related parasite *Toxoplasma gondii*, where TgMyoA is critical but not absolutely required for invasion (Meissner *et al*, 2002; Andenmatten *et al*, 2013; Bichet *et al*, 2016), so a greater understanding of the mechanical function of actomyosin during merozoite invasion is important. At least two energetic barriers during invasion require actomyosin activity, since chemical (Miller *et al*, 1979; Weiss *et al*, 2015) or genetic (Das *et al*, 2017; Perrin *et al*, 2018) disruption of actomyosin blocks the profound deformations of the RBC and merozoite internalisation. RBCs still undergo echinocytosis under these conditions suggesting that some breach of the RBC has still occurred. Biophysical modelling of RBC invasion suggests that actomyosin force may also be required for a third energetic barrier, to drive completion of entry and closure of the invasion pore (Dasgupta *et al*, 2014), however, no there is no evidence for this.

The importance of the biophysical properties of the RBC in determining the extent of any energetic barrier to parasite invasion has received increasing interest recently. Higher RBC membrane tension, for example, has been shown to reduce invasion success, whether due to natural variation or genetic polymorphisms such as the Dantu blood group (Kariuki *et al*, 2018). The *Plasmodium* merozoite appears to exploit the nature of these biophysical properties at multiple levels. For example, the binding of the parasite adhesin EBA-175 to its RBC receptor reduces the RBC membrane bending modulus (Koch *et al*, 2017; Sisquella *et al*, 2017). Overall, this paints a clear picture of invasion as an efficient balance of parasite force and modulation of the host cell biophysical properties (Dasgupta *et al*, 2014; Koch & Baum, 2016). However, a complete understanding of the energetic barriers found throughout the invasion process still remains unresolved.

Here, to gain insight into the energetics invasion and the role actomyosin activity plays during merozoite invasion, a conditional knockout approach was employed, building on the PfMyoA knockout (Robert-Paganin *et al*, 2019) to generate conditional mutations of PfMyoA and to target the auxiliary motor PfMyoB. PfMyosin B (MyoB) is a second *Plasmodium* class XIV localised to the extreme merozoite apex, suggesting a function during invasion, such as driving the initial stages of internalisation or organising apical organelles (Yusuf *et al*, 2015). Alongside a conditional knockout of light chain PfELC (Moussaoui *et al, manuscript submitted*), each mutant was analysed during merozoite invasion by video microscopy revealing three distinct stages of incomplete or aborted RBC invasion depending on actomyosin defect severity. The spectrum of phenotypes seen support the existence of three clear energetic barriers to successful entry, which together with previous works supports a stepwise model for actomyosin force action during merozoite invasion.

## Results

### Development of an ectopic expression platform for *Plasmodium* myosins

We demonstrated previously that PfMyoA is critical for asexual replication (Robert-Paganin *et al*, 2019). This was achieved using a conditional knockout system based on rapamycin (RAP)-dependent DiCre recombinase excision of the 3’ end of the *Pfmyoa* gene. Excision relies on *loxP* sites contained within synthetic introns (*loxPint*) (Jones *et al*, 2016) that were integrated into the *Pfmyoa* gene by selection-linked integration (SLI) (Birnbaum *et al*, 2017). As well as confirming that PfMyoA is critically important, we reasoned that this PfMyoA-cKO parasite line could be used as a platform for further investigation into the function of PfMyoA.

A strategy was developed to express mutant alleles of *Pfmyoa* from a second locus in the PfMyoA-cKO parasite line, to enable conditional mutation of any part of PfMyoA. We used the *p230p* locus, identified as dispensable throughout the parasite life cycle (van Dijk *et al*, 2010) and developed for targeted CRISPR/Cas9 integration (Ashdown *et al*, 2020, *in press*). To facilitate gene replacement on top of the PfMyoA-cKO background, the *p230p* targeting plasmid (pDC2-p230p-hDHFR) was modified by the exchange of *hdhfr* for *bsd*, which encodes the *blasticidin-S-deaminase* resistance gene (since PfMyoA-cKO parasites already express hDHFR) to form the pDC2-p230p-BSD targeting plasmid (Figure 1A). In the repair template plasmid (p230p-BIP-sfGFP, in which super-folder GFP (sfGFP) is inserted into the *p230p* locus) the constitutive BIP promoter was exchanged for the endogenous *Pfmyoa* promoter (prMA) for appropriate transgene expression timing (Figure 1A). The *Pfmyoa* promoter was amplified from 3D7 genomic DNA (defined as 2 kbp of sequence upstream of *Pfmyoa*) and introduced to form the p230p-prMA-sfGFP repair plasmid. The two plasmids were co-transfected into B11 (the parent line of PfMyoA-cKO) to generate p230p-prMA-sfGFP parasites, which exhibited merozoite-specific expression of sfGFP (Figure S1).

**Figure 1:**
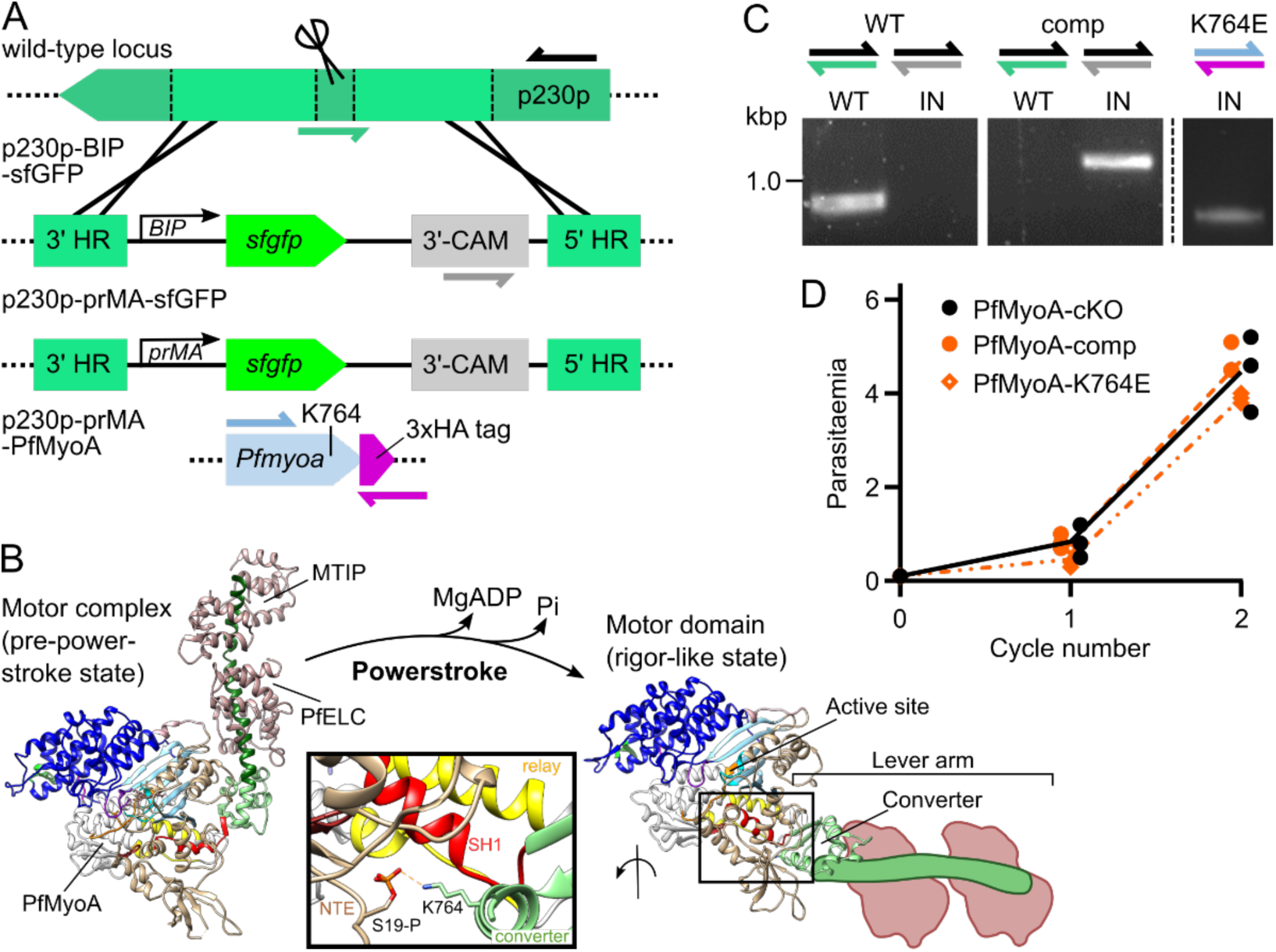
Development of a conditional knockout and complementation system for *Plasmodium* myosins. **A** Schematic of the modified *p230p* locus. The p230p-BIP-sfGFP repair template introduces *sfgfp* under a constitutive *BIP* promoter. The *BIP* promoter is exchanged for the *Pfmyoa* promoter, forming p230p-prMA-sfGFP, with *sfgfp* exchanged for 3xHA-tagged *Pfmyoa* forming PfMyoA-comp or PfMyoA-K764E. **B** Myosins produce force during a powerstroke, where conformational changes from ATP hydrolysis are communication by a relay (yellow) and SH1 (red) helices to the converter, to swing the lever arm. The SH1 helix in PfMyoA is unusually immobile and additional interactions are needed to stabilise the rigor-like state. Stabilising residues include those between phospho-S19 in the N-terminal extension (NTE, brown) and K764 in the converter (green). PPS structure from (Moussaoui *et al, manuscript submitted*); Rigor-like structure (PDB: 6I7D, neck region and light chain outlines added as schematic, by extension of the last helix of the converter). **C** Genotyping PCR of PfMyoA-comp and PfMyoA-K764E lines confirmed that the WT *p230p* locus (green half arrow) is completely lost in PfMyoA-comp, while the integrated locus (IN, grey half arrow) is present. **D** PfMyoA-comp parasites grow at the same rate as the parental PfMyoA-cKO line, while PfMyoA-K764E parasites grow slightly slower over 96 h. Lines show mean parasitaemia, N=3, each experiment in triplicate.

Having validated sfGFP expression in late stages, the p230p-prMA-sfGFP construct was further modified by exchange of *sfgfp* for *Pfmyoa* (re-codon optimised to avoid recombination) to form p230p-prMA-PfMyoA, which was transfected into the PfMyoA-cKO parasite line. In addition to wild type *Pfmyoa*, generating straight PfMyoA-complementation, an additional construct was made carrying a K764E mutation, forming PfMyoA-K764E parasites. The K764E mutation was designed to probe phospho-regulation of PfMyoA, wherein the charge reversal should repel phospho-S19 and prevent the stabilising effect of the K764-phospho-S19 interaction, proposed to enable fast cycling of PfMyoA in fast gliding stages (Robert-Paganin *et al*, 2019). This mutation should leave merozoites unaffected if they only need PfMyoA for maximal force production during invasion (Figure 1B). PfMyoA-comp and PfMyoA-K764E parasites were successfully generated, and the modification was confirmed by genotyping PCR, with no detectable WT remaining in PfMyoA-comp parasites (Figure 1C). Two independent attempts to generate corresponding mutations in S19 (testing the inverse site to K764), or to delete the entire N-terminal extension (residues 2-19) were unsuccessful.

Comparison of parasite growth over 96 h, without RAP induction, revealed no growth defect in PfMyoA-comp compared to the parental line (Figure 1D). The PfMyoA-K764E line demonstrated a 94% relative fitness per cycle (Figure 1D). This was unexpected, since the endogenous *Pfmyoa* locus is still present, suggesting that the second allele exhibits a slight dominant negative effect. This could explain the failure of transfection for more disruptive mutations in the N-terminal extension.

### Conditional complementation and mutagenesis of PfMyoA

To conditionally ablate the endogenous *Pfmyoa* allele, synchronised ring stage parasites were treated with rapamycin (RAP, 16 h, 100 nM), or DMSO as a control, in cycle 0 and parasitaemia was quantified by flow cytometry in each of the following three cycles. Excision of the *Pfmyoa* locus was verified by genotyping PCR and Western blot (Figure 2B-C). RAP-treated PfMyoA-comp parasites grew indistinguishably from DMSO-treated parasites, confirming that the severe growth defect seen in PfMyoA-cKO parasites is due to the truncation of PfMyoA. In contrast, RAP-treated PfMyoA-K764E parasites had a moderate growth defect, growing at, on average, 55% of DMSO-treated controls per cycle (Figure 2A).

**Figure 2:**
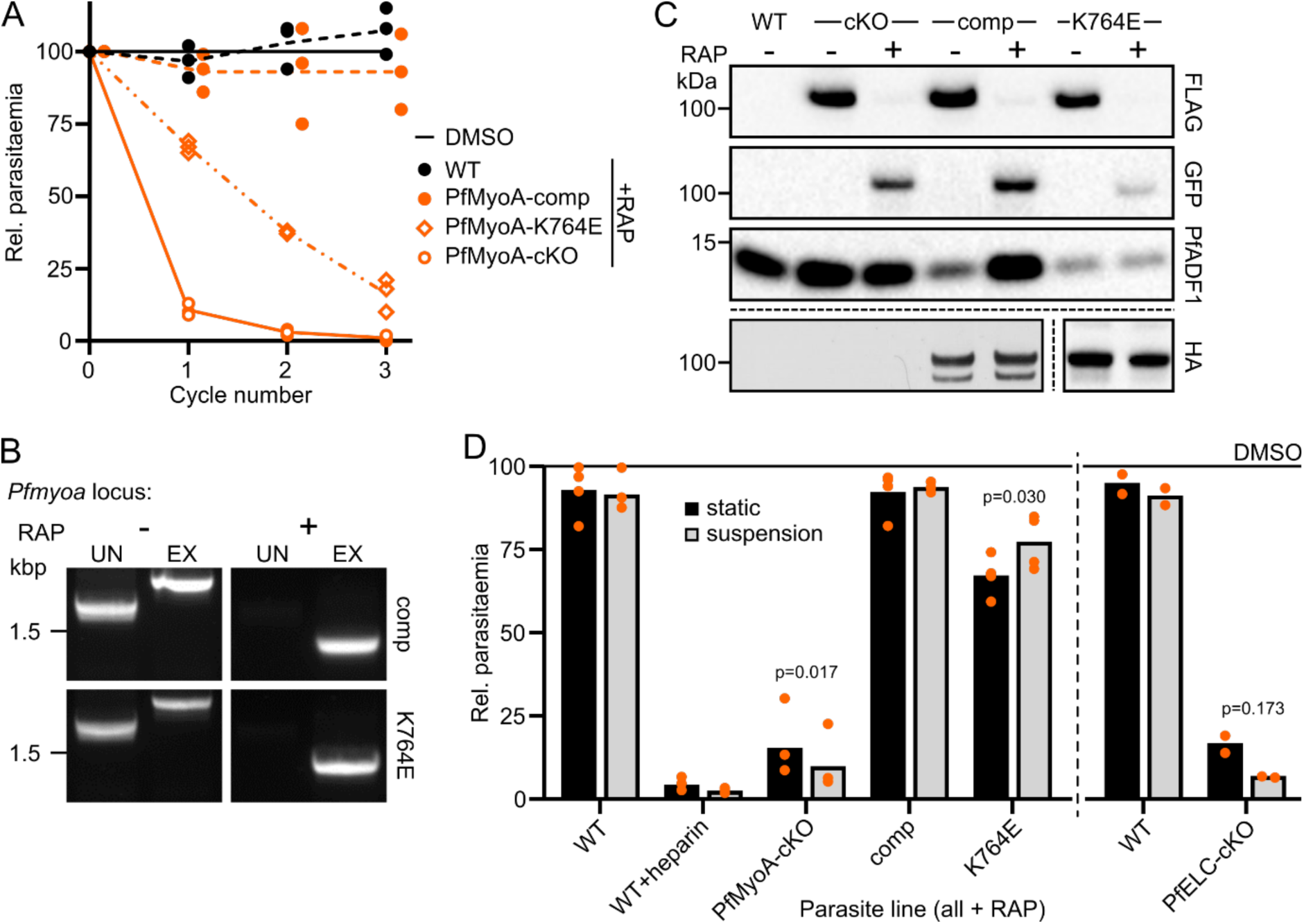
Conditional complementation and mutagenesis of PfMyoA. **A** RAP treatment of PfMyoA-comp and PfMyoA-K764E show that the complementing line has no growth defect, while the RAP-treated K764E line grows at around 55% of DMSO-treated control per cycle under static conditions. **B** Genotyping PCR of the endogenous *Pfmyoa* locus in PfMyoA-comp and PfMyoA-K764E after RAP treatment shows that the replacement of the unexcised, integrated allele (IN) with the excised allele (EX) is almost complete. **C** Western blot of WT, PfMyoA-cKO, PfMyoA-comp and PfMyoA-K764E schizonts confirms that PfMyoA-FLAG, expressed from the endogenous *Pfmyoa* locus, is almost completely lost in favour of truncated PfMyoA-GFP. PfMyoA-3xHA, expressed at the ectopic locus, is unaffected by RAP treatment. **D** Parasites were treated with DMSO or RAP and cultured under static or suspension conditions. Parasitaemia was then measured in the following cycle by flow cytometry and normalised to DMSO control for each line and condition, showing that suspension conditions partially alleviate the growth defect caused by K764E mutation, from 67% of DMSO-treated to 77%. Bars show mean parasitaemia, N=4 (or 2 for PfELC-cKO, tested separately), each experiment in triplicate. Significance assessed by paired t test, two tailed.

The PfMyoA light chain PfELC is essential for asexual replication (Moussaoui *et al, manuscript submitted*), but *in vitro* data shows that the complex of PfMyoA and MTIP can translocate actin without PfELC, albeit at half the speed (Bookwalter *et al*, 2017). This suggests that the absence of PfELC leaves a functional but strongly weakened motor. In light of the recent, unexpected demonstration that *P. falciparum* merozoites glide on a substrate when in static culture (Yahata *et al*, 2020), the static RAP growth assays were repeated, including PfELC-cKO, split equally between static and suspension conditions (Figure 2D). The strong replication defects in PfMyoA-cKO and PfELC-cKO lines were enhanced under suspension culture. In contrast, the replication defect of PfMyoA-K764E parasites was partially alleviated under suspension culture (from 67% of DMSO to 77%, p=0.03), consistent with the K764-phospho-S19 interaction being dispensable for invasion when gliding is bypassed by suspension culture (Figure 2D).

### Disruption of PfMyoB produces a mild growth defect

A contributor to the residual invasion seen in *T. gondii* MyoA-cKO parasites is myosin redundancy (Frénal *et al*, 2014). *Plasmodium* spp. lack the paralogues of TgMyoA, but do possess two genus-specific myosins, PfMyoB and PfMyoE, that could support PfMyoA during invasion. Genetic deletion of *P. berghei* MyoB caused no obvious defect throughout the life cycle (Wall *et al*, 2019), so to confirm whether PfMyoB is also dispensable and to investigate its role during invasion, a conditional KO was designed based on the PfMyoA-cKO line. Due to the difficulties in obtaining a pure transgenic population with the PfMyoA-cKO construct using SLI, a CRISPR-mediated strategy was developed for insertion of *loxPint* modules and a 3xHA-tag to the *Pfmyob* locus (Figure 3A). The construct was designed to conditionally excise 204 amino acids at the PfMyoB C-terminus, including the lever arm (containing the MLC-B light chain binding site) and part of the core motor domain, which on excision would form a truncated protein fused to sfGFP (Figure 3B). PfMyoB-cKO parasites were generated from the DiCre-expressing B11 line and verified by genotyping PCR (Figure 3C), showing no residual wild-type parasites. Culturing PfMyoB-cKO parasites alongside the parental line over 96 h in the absence of RAP showed no difference in growth (Figure 3D).

**Figure 3:**
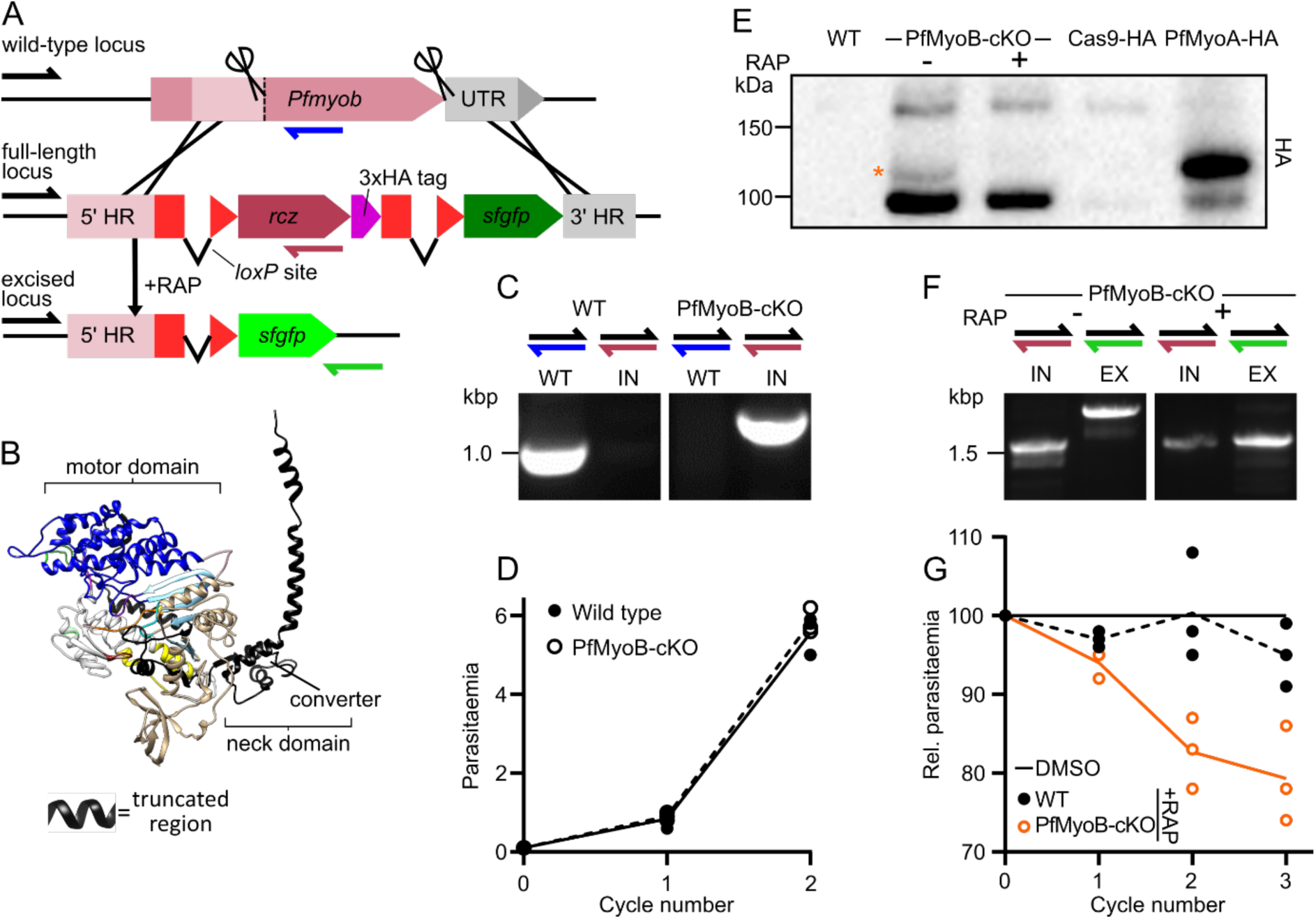
Disruption of PfMyoB produces a mild parasite growth defect. **A** Schematic showing generation of a PfMyoB-cKO targeting construct. A region of *Pfmyob* encoding the C-terminal 204 residues was synthesised with re-optimised codons (rcz) and a 3xHA tag, and is placed between two *loxPint* modules, with *sfgfp* out-of-frame downstream. Guide RNA sites (scissors) and homology regions were chosen to start as close as possible to the start and end of the modified region. **B** A structural model of PfMyoB indicating the region excised in PfMyoB-cKO (in black). **C** Genotyping PCR confirms that transfectants contain only the integrated locus (IN, purple half arrow), while the WT locus (blue half arrow) is completely lost. **D** Growth of PfMyoB-cKO parasites over 96 h is no different to the parental, DiCre-expressing, B11 line. Line shows mean parasitaemia, N=3, each experiment in triplicate. **E** Western blot analysis of WT, PfMyoB-cKO or Cas9-3xHA-expressing controls (where Cas9 was the only 3xHA-tagged protein or PfMyoA-3xHA was also expressed). In all lanes with PfMyoB-cKO or Cas9-3xHA-expressing controls a band around the expected size of Cas9-3xHA (168 kDa) and a presumed Cas9-3xHA breakdown product (∼95 kDa) is observed. In PfMyoB-cKO+DMSO, but not +RAP, a slightly larger band is detected around the expected size for PfMyoB-3xHA (97 kDa), confirming that PfMyoB-3xHA is properly expressed and lost after RAP treatment. The PfMyoB-3xHA band runs at a similar size to PfMyoA-3xHA control (96 kDa). **F** Genotyping PCR shows the loss of much of the integrated, unexcised locus (IN, purple half arrow) after RAP treatment and detection of the excised locus (EX, green half arrow). **G** Measuring the parasitaemia of PfMyoB-cKO parasites in each of the three cycles following RAP treatment shows a small, steady growth defect, of 93% on average. Lines show mean parasitaemia, normalised to DMSO for each line/cycle. N=3, each experiment in triplicate.

RAP treatment of PfMyoB-cKO parasites produced a small, consistent growth defect, with a fitness of 93% per cycle relative to DMSO treatment, compared to WT parasites which had a relative fitness of 98% per cycle (Figure 3G). Schizont samples taken at the end of cycle 0 and analysed by genotyping PCR or Western blot confirmed that excision was almost complete (Figure 3E,F). Therefore, disruption of PfMyoB produces a small growth defect, and if PfMyoB is involved in merozoite invasion, its function is not essential in the presence of functional PfMyoA.

### Video microscopy of merozoite invasion

Quantification of the growth defects observed in PfMyoA-cKO, PfMyoA-K764E, PfELC-cKO or PfMyoB-cKO gives only limited information about their cellular function. Video microscopy has long been used to describe merozoite invasion (Dvorak *et al*, 1975; Gilson & Crabb, 2009; Weiss *et al*, 2015) and more recently to uncover the effects of RBC and parasite mutants (Yap *et al*, 2014; Volz *et al*, 2016; Kariuki *et al*, 2018). However, the technique is not often used to capture large data sets on multiple parasite lines, due to the time required to capture and analyse the videos.

Having generated a panel of mutants addressing parasite myosin functions, we set out to analyse merozoite invasion by video microscopy after DMSO and RAP treatment. As in previous assays, parasites were treated, then incubated for ∼40 h until near the end of the same cycle. Purified schizonts were arrested before egress using PKG inhibitor ML10 (Baker *et al*, 2017) to increase synchronicity and schizont maturity. In turn, samples of each line were washed thoroughly to remove the drug and videos were captured in duplicate, for each of two independent biological repeats. Brightfield videos were recorded (3 fps, 10 minutes) and green fluorescence was captured at the start and end, permitting the exclusion of parasites that had not undergone proper excision after RAP treatment, except for PfELC-cKO which did not include a GFP-tag. Attempted invasion was defined as any merozoite attachment to an RBC that triggers echinocytosis. Almost 1300 invasion events were captured (between 88-204 for each line and treatment), of which around 55% were clear throughout the event, resulting in a dataset of 692 events.

Merozoite invasion can be broken down into attachment to the RBC, followed by deformation, internalisation and echinocytosis (Figure 4A). Depending on how many phases were achieved in an event, we classified invasion using a scheme adapted from (Yap *et al*, 2014) as either: (Type A) successful invasion; (Type B) internalisation incomplete and ejection of the merozoite; (Type C) deformation present but no internalisation; or (Type D) neither deformation or internalisation present, just attachment (Figure 4E). Comparison of the distribution of event types across the DMSO-treated lines shows that successful invasion was the most common event. Unexpectedly, DMSO-treated PfMyoB-cKO parasites had a much higher rate of invasion success (91% vs next highest 71%, p=0.038 PfMyoB-cKO vs others) (Figure 4C) suggesting that the other lines had slightly impaired invasion even after DMSO treatment, perhaps due to the sensitivity of PfMyoA or PfELC to epitope tags or the SLI machinery.

**Figure 4:**
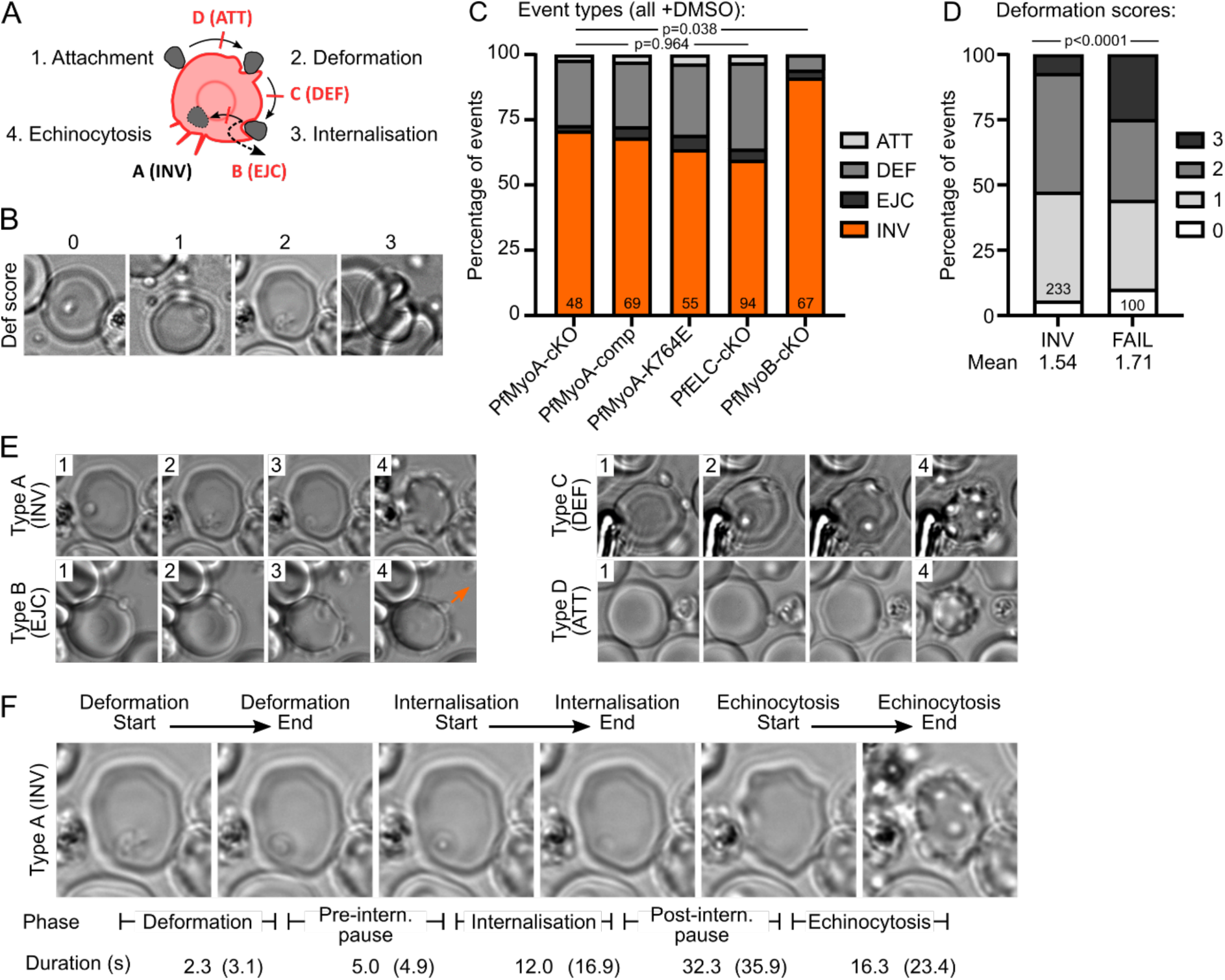
Video microscopy of merozoite invasion. **A** Schematic of merozoite invasion. Invasion comprises attachment to the RBC, deformation of the RBC membrane, internalisation of the merozoite and RBC echinocytosis. Invasion attempts are classified as successful (Type A) or by the phase of failure (Type B-D). **B** Each event was assigned a score based on the intensity of RBC deformation, from 0 (no deformation) to 3 (severe deformation). **C** Event types in videos from each line after DMSO treatment. While the distribution of events is not significantly different in PfMyoA or PfELC-cKO lines, PfMyoB-cKO shows significantly more successful invasion. Videos pooled from two independent experiments, each in duplicate. Numbers indicate total videos. Significance assessed by chi-square test, either each PfMyoA/PfELC line separately, or PfMyoB-cKO vs others pooled. **D** Deformation scores from all DMSO-treated lines separated by successful invasion (INV) or any type of failure (FAIL) reveal a significant increase in strong deformation in failed events. Numbers indicate total videos. Significance assessed by chi-square test. **E** Examples of each event type, with the numbers indicating the phase of invasion shown in that image (numbers from **A**). (For Type B event, see Video S6; for Type D event, see Video S4). **F** For Type A and Type B events, the start and end of deformation, internalisation and echinocytosis were timed, leading to the five intervals shown. The median times from Type A events across all lines after DMSO treatment is shown, with overall agreement to published values ((Gilson & Crabb, 2009) in brackets).

A qualitative score was assigned to each event based on the intensity of the deformation, from 0 (no deformation, just attachment) to 3 (severe deformation of the RBC extending across the cell), developed by (Weiss *et al*, 2015) (Figure 4B). Comparison of the deformation scores between successful invasion events and invasion failures shows that invasion failures had significantly stronger deformations (p<0.0001) with a higher mean deformation score of 1.71, compared to 1.54 (Figure 4D).

For successful invasion and Type B failures, the scheme of (Gilson & Crabb, 2009) was adapted for timing the duration of five phases: deformation, internalisation and echinocytosis and the two pauses: before internalisation, when the TJ is thought to be formed, and after internalisation, before echinocytosis. Comparison of the phase timings from DMSO-treated parasites with published values shows similar results (Gilson & Crabb, 2009) (Figure 4F). Since the events for each line were pooled across two biological repeats, an example comparison was made between the two biological repeats for PfMyoA-comp after DMSO treatment (Figure S2), confirming that they were highly similar.

Type B failures as defined previously (Yap *et al*, 2014) (there called Type III invasion) as merozoites that did not produce a ring despite triggering echinocytosis, due to failure of resealing. This was sometimes followed by ejection of the merozoite to the outside of the RBC. In our video observations, Type B failures occurred in 4% of DMSO-treated parasite events (13/333) and were defined as any event where ejection of merozoites to the outside of the RBC was observed. In Type B failures the invasion attempt consistently begins normally and only after echinocytosis is complete is the merozoite ejected from the RBC through the same invasion pore. Compared to Type A invasion, Type B invasion attempts had slower internalisation and a trend towards a longer pause before internalisation (Figure S3).

Ejection presumably occurs due to a defect in resealing the pore, as it was only observed long after the apparent end of internalisation, though the driving force behind the ejection remains unclear. In some videos the invasion pore remained visible until ejection of the merozoite, while in others the merozoite appeared motile throughout, performing a swirling motion during and after ejection (Figure S3, Video S6). After ejection, merozoites typically remained attached to the RBC and none underwent a second invasion attempt before the end of the video. Type B failures may result from a combination of lower parasite force production and increased RBC biophysical resistance, as described by biophysical models of invasion, which includes the role of the TJ in providing line tension to close the pore (Dasgupta *et al*, 2014).

### Without PfMyoA or PfELC, merozoites cannot strongly deform or internalise

Conditional KO of PfMyoA showed an almost complete growth defect (Robert-Paganin *et al*, 2019) and accordingly, PfMyoA-cKO parasites after RAP treatment showed zero successful invasion events (0/53) (Figure 5A). In all events recorded, PfMyoA-cKO + RAP showed no deformation or internalisation (Type D failure, Video S4). Since there was a complete block at deformation, this mutant cannot be used to probe the role of the motor in internalisation directly. For PfMyoA-comp parasites, the event types observed in DMSO- and RAP-treated parasites were slightly different, with a moderate drop in invasion success (from 68% to 52%) and corresponding increase in Type C failures (p=0.047) (Figure 5A). However, there were no significant differences in deformation strength or phase timings suggesting that overall the PfMyoA-comp protein successfully complemented the function of the native protein (Figure 5B-C).

**Figure 5:**
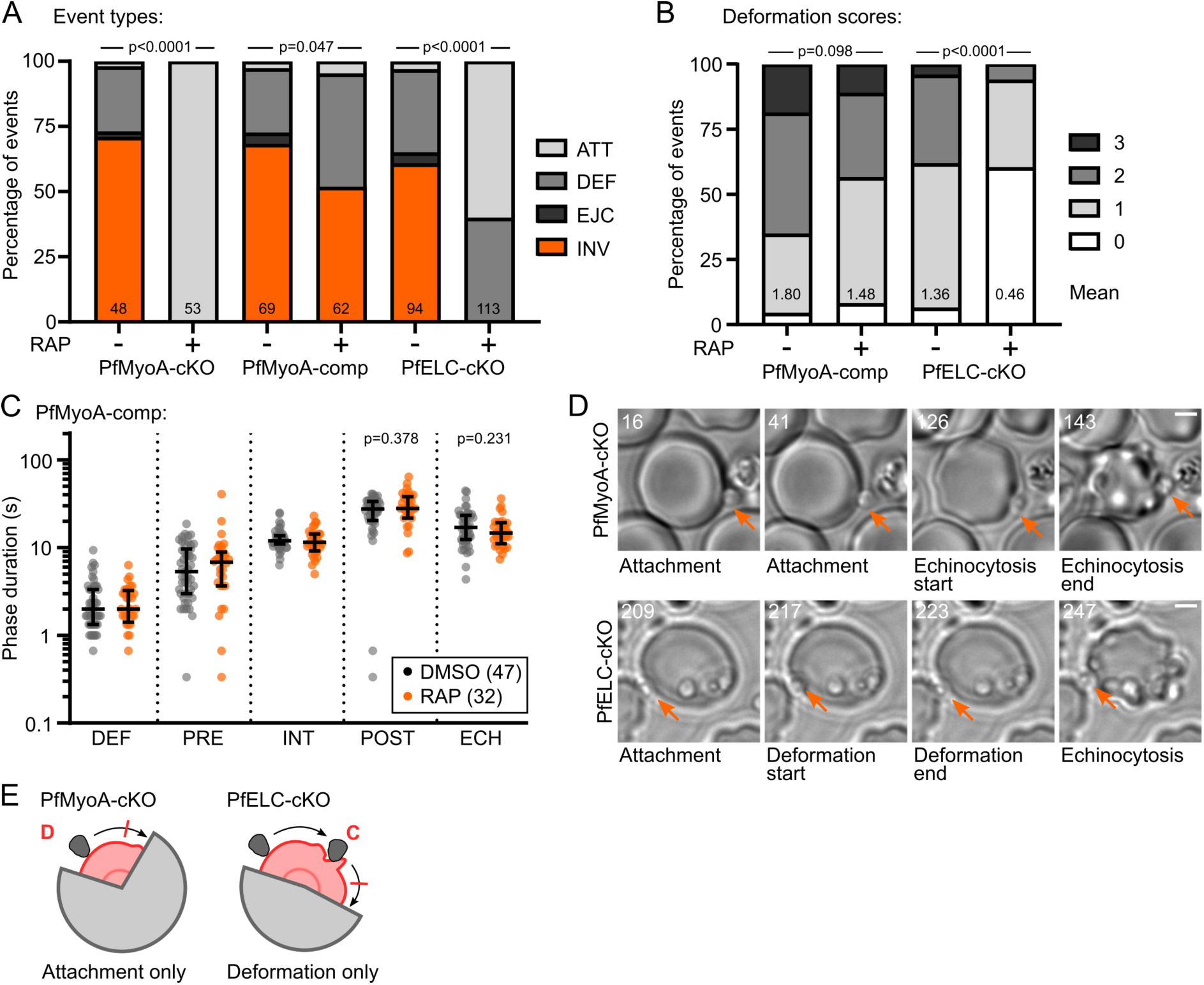
In the absence of PfMyoA or PfELC, merozoites cannot strongly deform or internalize. **A** Comparison of event types from PfMyoA-cKO, PfMyoA-comp and PfELC-cKO lines after DMSO and RAP treatment. PfMyoA-cKO parasites show neither deformation nor internalisation after RAP treatment (p<0.0001, significance assessed by Fisher’ s exact test comparing pooled failures to Type A events). In contrast, PfMyoA-comp shows only a slight drop in the rate of successful invasion after RAP treatment (p=0.047). PfELC-cKO parasites also shows a complete loss of successful invasion, but almost half of the events did involve deformation. Significance assessed by chi-square test. **B** The distribution of deformation scores does not change significantly for PfMyoA-comp events after RAP treatment. The mean deformation score for PfELC-cKO merozoites is greatly reduced by RAP treatment. Significance assessed by chi-square test. **C** Comparing the duration of each phase of invasion for PfMyoA-comp parasites after DMSO or RAP treatment shows no significant differences. **D** Examples of RAP-treated PfMyoA-cKO merozoite, showing attachment only, with no further progress (Video S4), and a PfELC-cKO merozoite undergoing a Type C failure, showing deformation but no internalisation (Video S5). Time indicated in seconds, scale bar 2 μm. **E** Schematic based on Figure 4A showing invasion attempts by PfMyoA-cKO merozoites arrest after attachment, while invasion attempts by PfELC-cKO merozoites arrest after deformation, though many arrest before deformation as well.

Having confirmed the importance of PfMyoA for merozoite force production, we next asked whether the weakened motor present in PfELC-cKO might reveal more about the phases of invasion that require actomyosin force. Like PfMyoA-cKO parasites, RAP-treated PfELC-cKO parasites did not achieve any successful invasions (0/113) (Figure 5A). However, 40% of PfELC-cKO events were able to deform the RBC (Type C events, Video S5). Though the deformation scores were significantly weaker (p<0.0001, mean shifted from 1.36 to 0.46), this shows that a partially functional motor can achieve inefficient deformation (Figure 6B). Importantly, no PfELC-cKO merozoites were able to initiate internalisation, supporting a critical role for PfMyoA in driving merozoite internalisation, as well as deformation, and suggesting that the process of internalisation has a higher energetic barrier than deformation.

**Figure 6:**
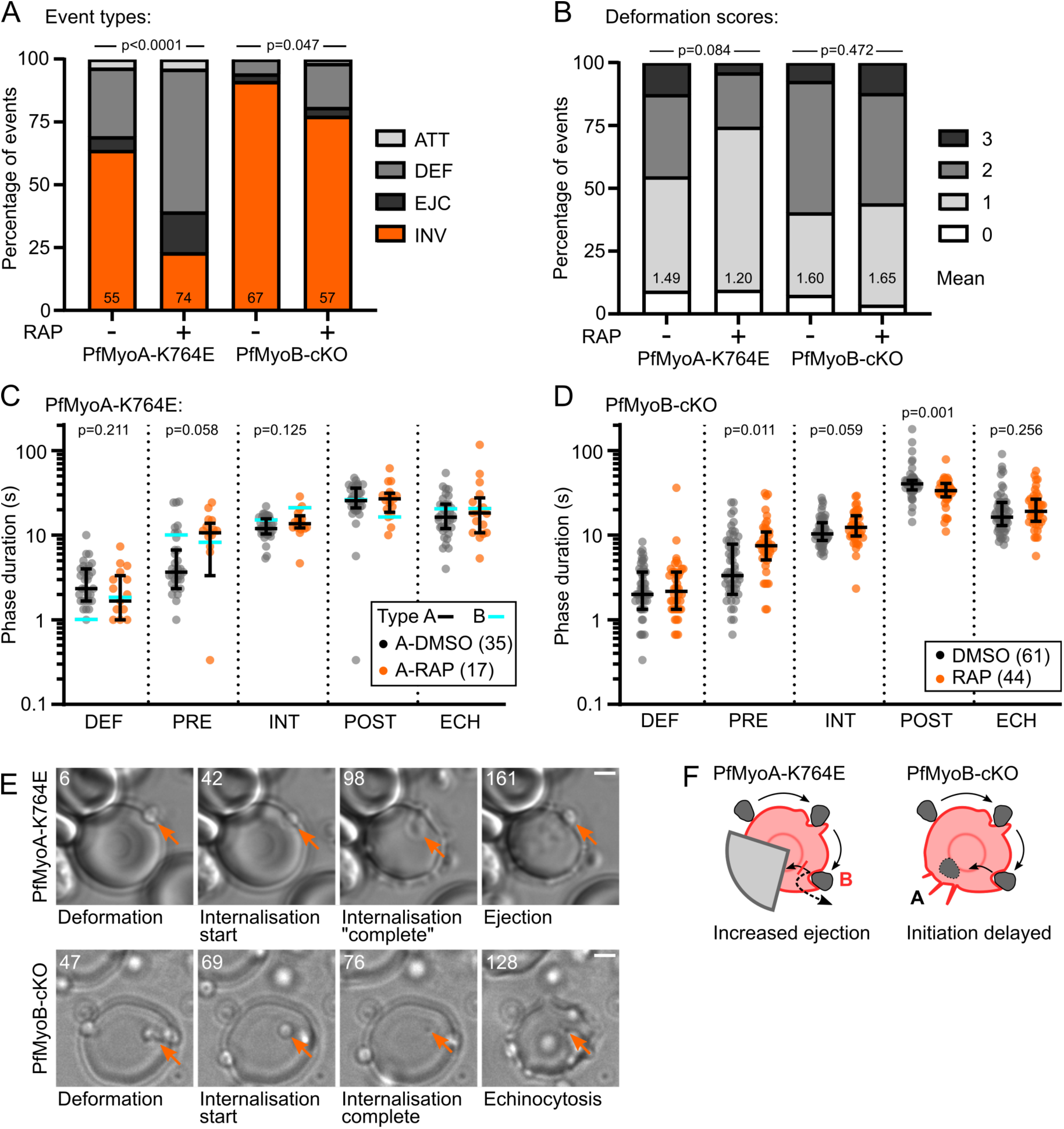
PfMyoA drives invasion pore closure, while PfMyoA and PfMyoB both help initiation of internalization. **A** RAP treated PfMyoA-K764E parasites showed a significantly shifted distribution of event types, with more Type B and Type C events. This is consistent with these parasites having insufficient motor function to overcome a third barrier, at completion of internalisation. Significance assessed by chi-square test. PfMyoB-cKO parasites showed a significant increase in Type C failures, consistent with an impairment at initiation of internalisation. Significance assessed by Fisher’ s exact test, comparing successful invasion to pooled invasion failures. **B** PfMyoA-K764E parasites show a slight weakening of deformation, though not significant. There is no difference between the deformation scores in PfMyoB-cKO parasites after RAP treatment. Significance assessed by chi-square test. **C** For PfMyoA-K764E parasites undergoing Type A events (black bars and data points) RAP treatment induces a longer pause pre-internalisation. Only in Type B events (cyan bars) after RAP treatment is internalisation significantly slower and the pause post-internalisation shorter. Bars show median and interquartile range, or median only for Type B events. Significance assessed between Type A DMSO and RAP treatments by Mann-Whitney test, shown when p<0.5. **D** The duration of pre-internalisation pause is significantly increased in PfMyoB-cKO parasites, suggesting that PfMyoB plays a role in initiation of internalisation. The post-internalisation pause is reduced by a similar amount. Bars show median and interquartile range. Significance assessed by Mann-Whitney test, shown when p<0.5. **E** Example of a RAP-treated PfMyoA-K764E merozoite, undergoing a Type B failure, showing apparent completion of internalisation before subsequent ejection (Video S6), and a PfMyoB-cKO merozoite undergoing successful invasion (Video S7). Time indicated in seconds, scale bar 2 μm. **F** Schematic based on Figure 4A showing that PfMyoA-K764E merozoites can proceed to internalisation, but are frequently ejected, while PfMyoB-cKO merozoites invade successfully, but with delayed initiation of internalisation.

### PfMyoA drives invasion pore closure, while PfMyoA and PfMyoB both help initiation of internalization

Understanding the role of motor force during internalisation depends on finding intermediate-strength motor mutants able to initiate internalisation. Alone amongst the PfMyoA mutants PfMyoA-K764E merozoites could initiate invasion, though less efficiently, showing a marked increase in Type B failures (Video S6), from 5% to 16%, as part of a significant disruption to event types (p<0.0001, Figure 6A). Deformation was not significantly affected, but there was a notable (though only borderline significant) increase in the median duration of the pre-internalisation pause after RAP treatment, from 3.7 s to 10.7 s in Type A events (Figure 6C, p=0.058). In Type B events, this much longer pre-internalisation pause was also present, but in both DMSO- and RAP-treated parasites. This suggests that either a weaker motor or a more resistant RBC can delay the initiation of internalisation.

In contrast, internalisation itself was only significantly slowed in RAP-treated PfMyoA-K764E parasites undergoing Type B events, not Type A events (Figure 6C) (21 s, compared to 13.7 s for Type A events, p=0.003). Internalisation in DMSO-treated parasites undergoing Type B events was slightly slower than Type A (15 s vs 12 s), but to a much lesser extent. Therefore, only the combination of a weaker motor and a more resistant RBC resulted in strongly slowed internalisation. This may reflect slower motion during internalisation or an arrest at completion of internalisation.

Therefore, following RAP treatment, PfMyoA-K764E parasites are more likely to fail at initiation of internalisation (an increase in Type C failures, Figure 6A) and, when they can initiate it, they take longer to do so (a longer pre-internalisation pause, Figure 6C). Importantly, these parasites are also more likely to fail to complete internalisation (causing the increase in Type B failures, Figure 6A) and they take much longer to internalise when they fail, and slightly longer even when successful (Figure 6C).

Having demonstrated the effect of a gradient of PfMyoA motor defects in invasion, we finally sought to test the role of PfMyoB in the invasion process. Video microscopy of RAP-treated PfMyoB-cKO parasites showed only a moderate reduction in successful invasion (from 91% to 77%, p=0.045, Video S7) and increase in Type C failures (from 6.0% to 17.5%) (Figure 6A), while the distribution of deformation scores was unchanged (p=0.472) (Figure 6B). The durations of some invasion phases were affected by PfMyoB-cKO. The pre-internalisation pause was more than doubled (from 3.3 s to 7.5 s, p=0.011) and the pause post-internalisation was significantly reduced (from 40.3 s to 33.7 s, p=0.001) by an amount roughly equal to the combined increases in duration of earlier phases (Figure 6C).

While the overall defect in PfMyoB-cKO parasites was mild (a moderate increase in Type C failures), the significant delay in initiation of internalisation is consistent with a model of PfMyoB supporting the first stages of translocating the TJ. However, this role of PfMyoB, or any other, is clearly not required for internalisation. The shorter pause post-internalisation directly corresponds to the delays earlier in invasion, suggesting that the onset of echinocytosis falls at a set time after the stimulus regardless of the timing of subsequent phases, an effect also seen in PfMyoA-K764E parasites and in a previous study (Weiss *et al*, 2015).

Therefore, while PfMyoB-cKO merozoites are delayed in initiation of internalisation, and PfMyoA-cKO and PfELC-cKO merozoites have insufficient force production to overcome the steps of deformation or internalisation, PfMyoA-K764E merozoites show a distinct defect at a third energetic barrier: completion of internalisation. These data therefore support a three-step model for the energetics of red blood cell entry by the merozoite: surface deformation; initiation of internalisation/entry; and completion of internationalisation/closure of the tight junction.

## Discussion

### PfMyoA-K764E moderately impairs invasion and may block gliding

By extending the conditional knockout platform developed for PfMyoA (Robert-Paganin *et al*, 2019) to the auxiliary motor PfMyoB, the essential light chain PfELC and combining this with conditional complementation of PfMyoA, we have generated a series of malaria parasite motor mutants that show a range of defects in their ability to enter red blood cells. By investigating these defects by video microscopy we have revealed three different energetic barriers to invasion, each requiring some level of motor activity (Figure 7A).

**Figure 7:**
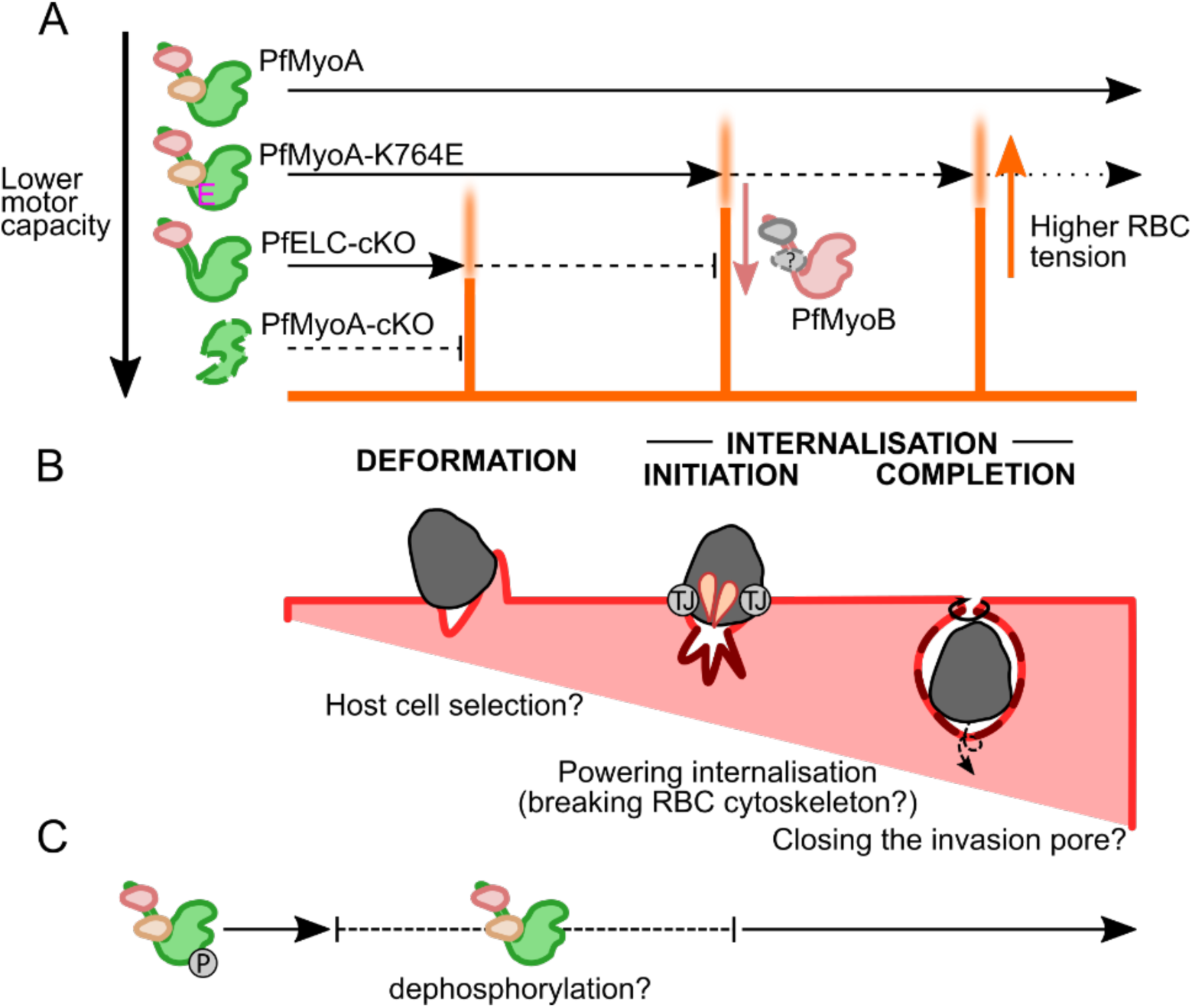
A stepwise model for *Plasmodium* myosin force generation during merozoite invasion. **A** PfMyoA produces force at three sequential energetic barriers to drive invasion. Successively more disruptive mutations bring the PfMyoA force production capacity closer to, or below, the energetic barriers, represented by orange bars with blurred tops indicating variability in individual parasites and RBCs. Some PfMyoA-K764E parasites fail at initiation of internalisation (Type C failures), some fail at completion of internalisation (Type B failures) and relatively few complete invasion. PfELC-cKO parasites often fail at the barrier of deformation (Type D failures), though some can deform and fail at initiation of internalisation (Type C failures). PfMyoA-cKO parasites never overcome the energetic barriers to deformation or internalisation. PfMyoB-cKO parasites are only impaired at initiation of internalisation, suggesting that PfMyoB may act to reduce the energetic barrier at initiation of internalisation. **B** Overview of actomyosin functions at each of the three steps, including using deformability to select suitable host cells, driving initial wrapping of the merozoite and twisting shut the invasion pore. Parasite effectors reduce the energetic barriers by modulating RBC biophysical properties, including creation of the tight junction (TJ) to introduce a line tension, insertion of membrane material (darker membrane) and binding of adhesins like EBA-175 that may affect RBC lipid packing **C** PfMyoA is phosphorylated before egress and may be dynamically dephosphorylated at some point before internalisation to tune the motor for maximal force production.

For flexible expression of conditional *Pfmyoa* mutations, a distal expression site was developed at the *p230p* locus, dispensable for development throughout the life cycle (van Dijk *et al*, 2010). PfMyoA-complementation and PfMyoA-K764E alleles were successfully integrated into the *p230p* locus, but other mutants affecting the *Pfmyoa* N-terminus, which have an equal or stronger impact on PfMyoA function *in vitro* (Robert-Paganin *et al*, 2019), could not be generated after two transfection attempts. Expression of a second copy of *Tgmyoa* produces a strong down-regulation at the endogenous locus (Hettmann *et al*, 2000; Meissner *et al*, 2002; Herm-Götz *et al*, 2007), which may explain the failure of transfections with moderately or strongly defective *Pfmyoa* alleles.

PfMyoA-K764E was confirmed to have a moderate phenotype after RAP treatment under static conditions while suspension conditions alleviated around 40% of the defect, consistent with the hypothesis that the interaction between phospho-S19 and K764 is critical only in stages where gliding is required (Robert-Paganin *et al*, 2019). The culture conditions that best imitate physiological conditions are not clear, but suspension culture significantly affects invasion phenotypes and adhesin expression (Paul *et al*, 2015; Awandare *et al*, 2018; Nyarko *et al*, 2020). The recent demonstration that merozoites exhibit actin-dependent gliding (Yahata *et al*, 2020) might explain why PfMyoA-K764E had a stronger effect on static invasion, if merozoites first tune PfMyoA for gliding before S19 is dephosphorylated to tune the motor for invasion (Figure 7C). The extensive calcium ion signalling that regulates organelle secretion during merozoite invasion might modulate PfMyoA phosphorylation, since a calcium-dependent phosphatase, calcineurin, critically regulates merozoite attachment (Paul *et al*, 2015; Philip & Waters, 2015). To assess the broader phospho-tuning hypothesis will require mutation of PfMyoA-S19 in merozoites and the fast gliding sporozoite, as well as direct assessment of the phosphorylation state of PfMyoA-S19 at each phase of invasion.

### Deformation is the first step during invasion requiring PfMyoA force

By filming highly synchronised schizonts at high parasitaemia (∼50%) and selecting only events that result in echinocytosis, large numbers of events can be captured by video microscopy. Though many events where a merozoite attached but failed to form a TJ may be missed by this approach, around 75% of successful invasion events result in echinocytosis (Weiss *et al*, 2015), so a minority of successful invasion events will be missed. Since the focus of this study is on the later phases of invasion involving motor activity, this approach was deemed an acceptable compromise for the detection of a greater number of events.

Previous studies using chemical (Miller *et al*, 1979; Weiss *et al*, 2015) or genetic (Das *et al*, 2017; Perrin *et al*, 2018) inhibition of merozoite actomyosin have clearly shown that, without force production, merozoites cannot deform the RBC or begin internalisation. Biophysical modelling work has suggested that internalisation should present a third energetic barrier: transition from a “partially-wrapped” to “completely-wrapped” state at completion of internalisation (Dasgupta *et al*, 2014). However, the role of the actomyosin motor during internalisation has not been probed directly, due to the need for intermediate strength mutants that can overcome the earlier energetic barriers.

Consistent with these studies, PfMyoA-cKO merozoites were completely blocked in both deformation and internalisation. These two phases were almost completely restored in PfMyoA-comp parasites, confirming that these two energetic barriers require PfMyoA activity. As an aside, the slight fall in PfMyoA-comp invasion success in general may result from the use of altered codons or the absence of regulatory DNA sequences found beyond the 2 kb promoter sequence or in the two short *Pfmyoa* introns, omitted in the complementing allele.

Unlike PfMyoA-cKO, PfELC-cKO only reduced deformation, though it also blocked all internalisation, suggesting that the level of motor activity retained in PfELC-cKO parasites is very low, and that the energetic barriers downstream of deformation are higher. This agrees with the almost complete replication defect seen in PfELC-cKO parasites (Moussaoui *et al, manuscript submitted*).

Deformation may help select a RBC with suitable biophysical properties for invasion (Weiss *et al*, 2015), and the completion of this selection process could act as a checkpoint, triggering signalling for secretion of TJ components, and for the proposed dephosphorylation of PfMyoA. It was previously reported that invasion failure correlated with weaker deformation (Weiss *et al*, 2015). However, in the current study invasion failure correlated with stronger deformation. This is likely due to the exclusion of weaker merozoite contacts that did not lead to echinocytosis.

### The second step, initiation of internalisation, is facilitated by both PfMyoA and PfMyoB

Across the different mutants, a longer pause before internalisation correlated with reduced invasion success. For PfMyoB-cKO, this delay was the most striking defect during invasion, suggesting that the initiation of internalisation is when PfMyoB plays its role. Though unrelated to PfMyoB in sequence, *T. gondii* MyoH shares the extreme apical localisation and is required for initial translocation of the TJ over the parasite apex, “handing over” to the TgMyoA motor complex (Graindorge *et al*, 2016). PfMyoB could play a similar role to TgMyoH, though PfMyoB is not critical for invasion. Since PfMyoB was previously shown not to co-localise with the TJ during internalisation but instead stayed at the merozoite apex (Yusuf *et al*, 2015), its functional role might be indirectly related to internalisation, either by contributing to the stability of the merozoite apex or the secretion of invasion ligands.

The fall in successful invasion in PfMyoB-cKO parasites, from 91% to 77%, was roughly consistent with the mild growth defect, with relative fitness falling to 93% per cycle and consistent with work showing that constitutive knockout of *P. berghei* MyoB has no defect throughout the life cycle (Wall *et al*, 2019). Further insight into the function of PfMyoB may come from study of its light chains and other interaction partners. MLC-B, the one currently identified light chain, is very large like the PfMyoA regulatory light chain, MTIP. Unlike PbMyoB, PbMLC-B was resistant to knockout, so may perform an important structural function (Wall *et al*, 2019).

Overall, the increased pause before internalisation appears to be a symptom of a weaker motor, confirming that initiation of internalisation is the second energetic barrier that requires PfMyoA motor activity, supported by PfMyoB motor activity. This seems to be the most common energetic barrier for merozoites to fail at, since Type C failures are by far the most common in DMSO-treated parasites.

### Completion of internalisation is a third and final motor-dependent step

For the first time, we demonstrate a third energetic barrier at the completion of internalisation. In PfMyoA-K764E merozoites there was a trend towards slower internalisation and a striking, three-fold increase in the rate of merozoite ejection after apparently completing internalisation, suggesting that PfMyoA force production is required through to the end of invasion, for completion of internalisation.

Type B failures were identified in *P. falciparum* in AMA1-cKO parasites (Yap *et al*, 2014) where this phenotype was suggested to result from failure to reseal the RBC membrane after entry. A comparable ejection of *T. gondii* tachyzoites was observed after depletion of cAMP-dependent kinase PKA, referred to as “premature egress” (Uboldi *et al*, 2018) although the same behaviour was not reported in a PKA-cKO line in *P. falciparum* (Patel *et al*, 2019). Translocation of the nucleus through the narrow TJ with the help of a posterior pool of actin was proposed to be a limiting factor for internalisation of *T. gondii* tachyzoites, with acto-myosin mutants pausing mid-internalisation at the point of nuclear entry (Del Rosario *et al*, 2019). However, we did not observe a similar pause during merozoite internalisation, instead the defect observed in motor-impaired PfMyoA-K764E parasites came later, after apparent completion of internalisation, so translocation of the nucleus is unlikely to depend on PfMyoA or PfMyoB. Instead, *P. falciparum* Myosin E (MyoE) might perform this role, since it was identified in a nucleus-associated proteome (Oehring *et al*, 2012) and *P. berghei* MyoE localises to the merozoite posterior (Wall *et al*, 2019). PbMyoE interacts with multiple members of the PbMyoA motor complex (Fang *et al*, 2018) and genetic deletion of PbMyoE impaired sporozoite entry to the mosquito salivary glands (Wall *et al*, 2019), consistent with PfMyoE supporting PfMyoA during invasion. Future experiments targeting this motor specifically will be required to define its precise function.

As predicted by biophysical modelling (Dasgupta *et al*, 2014) PfMyoA appears to support closure of the invasion pore behind the merozoite. Careful observation of *T. gondii* tachyzoites at the completion of internalisation revealed that twisting motility was required for efficient closure of invasion pore (Pavlou *et al*, 2018). This twisting could also be employed by *Plasmodium* merozoites. A recent study demonstrated helical motility of *P. knowlesi* merozoites on a substrate (Yahata *et al*, 2020). In a similar fashion, we observed a “swirling” motion by *P. falciparum* merozoites both before internalisation and after ejection (Videos S6, S8). Though the twisting motility in tachyzoites was not TgMyoA-dependent (Pavlou *et al*, 2018) our observations of PfMyoA-K764E merozoites suggest that PfMyoA has a role in driving completion of internalisation, possibly by driving closure of the invasion pore.

In summary, this study has used conditional knockouts to dissect the roles played by parasite myosin motors during the process of red blood cell invasion and developed a model of three sequential energetic barriers that require active actomyosin force. Successively stronger defects in the PfMyoA motor complex point to roles for the motor at deformation and at initiation and completion of internalisation. Meanwhile, though PfMyoB is clearly not critical for invasion, it seems to support timely initiation of internalisation. Future work will be required to understand the effect of variation in red blood cell biophysical properties, the parasite effectors that manipulate RBC properties and how motor force is regulated to rapidly and efficiently power the steps of merozoite invasion, the first stage in the development of malaria pathogenesis.

## Acknowledgements

This work was funded by Wellcome through an Investigator Award to J.B. (100993/Z/13/Z), the Human Frontier Science Program (RGY0066/2016 to J.B.) and a PhD studentship to T.C.A.B. (109007/Z/15/A). We thank Anne Houdusse, Dihia Moussaoui and Julien Robert-Paganin for sharing pre-publication structures for the complete PfMyoA complex. We thank Kathrin Witmer for help with design of constructs. We thank Mike Blackman for generous provision of the B11 line, David Baker for provision of the ML10 inhibitor and Marcus Lee for provision of CRISPR/Cas9 plasmids.

## Author Contributions

All authors were involved in conceptualization and writing of the manuscript. T.C.A.B. and S.H. conducted experiments. T.C.A.B. carried out formal analysis. J.B. supervised and acquired funding for the project.

## Methods

### Software for DNA sequence analysis and protein structure prediction

DNA constructs were designed using Benchling (benchling.com) and guide RNAs using CHOPCHOP (Labun *et al*, 2019). Protein structure predictions were generated using Phyre2 (Kelley *et al*, 2015) and models were visualised using UCSF Chimera (Pettersen *et al*, 2004).

### DNA manipulation

PCR was carried out according to the manufacturer’s protocols using Phusion polymerase (NEB), or Advantage 2 Polymerase mix (Takara Bio) for amplification of UTRs and *Pfmyob*, and constructs were assembled by Gibson assembly with DNA inserts in a 1:3 molar ratio. For transfections, plasmids were purified from 100 ml of overnight culture using a plasmid maxiprep kit (Qiagen). Before transfection, plasmids were purified by ethanol precipitation (with 0.1 vol 3M sodium acetate pH 5.2, 1.5 vol 100% ethanol), washed in 70% ethanol, air dried and resuspended in TE buffer (10 mM Tris-HCl, 1 mM EDTA, pH 8.0).

For modification of the *p230p* locus in the PfMyoA-cKO background, targeting and repair constructs were modified from (Ashdown et al, *in press*). The targeting construct, pDC2-p230p-hDHFR, (originally adapted from (White *et al*, 2019)) carries 3xHA-tagged Cas9 and the *p230p*-targeting guide RNA. This was modified by excising *hdhfr* with NcoI/SacII and ligating in *bsd* to form pDC2-p230p-BSD. The *bsd* sequence was amplified from pB-CBHALO (Stortz *et al*, 2019), with the N-terminal sequence modified to MAK during amplification, to match a consensus sequence (Mesén-Ramírez *et al*, 2016). The p230p-BIP-sfGFP repair plasmid (pkiwi003, Ashdown et al, *in press*) was modified to excise the BIP promoter by SacII/NheI digestion, and ligation of a 2.0 kbp region directly upstream of *Pfmyoa* (roughly 2/3 of the intergenic region) to form p230p-prMA-sfGFP. The *Pfmyoa* cds with 3xHA-tag was generated synthetically (GeneART) with altered codons and ligated into p230p-prMA-sfGFP after NheI/PstI digestion to form p230p-prMA-PfMyoA. K764E, S19A or ΔN mutants were generated by site-directed mutagenesis.

PfMyoB-cKO was generated using the same two plasmid CRISPR/Cas9 system, with guide RNAs chosen to target the start of each of the upstream and downstream homology regions. pDC2-cam-co.Cas9-U6.2-hDHFR (White *et al*, 2019) was digested with BbsI and annealed guide RNA oligonucleotides were ligated into the pDC2 backbone, forming pDC2-PfMyoB-hDHFR-1 or -2. The repair plasmid was designed like the PfMyoA-cKO construct (Robert-Paganin *et al*, 2019), but without the SLI machinery and constructed from a generic pUC19 backbone. The *loxPint* modules flanked the 611 bp codon-optimised region (rcz) and 3xHA tag, and this was synthesised with the downstream *sfgfp* (GeneART) and assembled with upstream and downstream homology regions of 671 bp and 881 bp amplified from genomic DNA. The sites targeted by the two guide RNAs were altered in the rcz region, so no additional shield mutations were required.

### Parasite culture and transfection

*P. falciparum* strains B11 (Perrin *et al*, 2018) and PfMyoA-cKO (Robert-Paganin *et al*, 2019) were cultured in complete culture media (CCM) comprising RPMI 1640 (Life Technologies) supplemented with 0.5% w/v Albumax-II (Gibco) under standard conditions (Trager & Jensen, 1976). Parasites were cultured at 4% haematocrit (using human O+ RBCs) and synchronised with 5% sorbitol (Sigma). For transfection, parasites were grown to 5% at ring-stage and electroporated with 50 µg of each of the targeting and repair plasmids (or 25 µg each for the two targeting plasmids for PfMyoB-cKO). Purified plasmids were resuspended in a total volume of 50 µl of TE buffer (pH 8.0) added to 350 µl sterile cytomix buffer (Adjalley *et al*, 2010). Plasmid uptake was selected for by adding fresh 2.5 nM WR99210 (Jacobus Pharmaceutical) or 5 µg/ml blasticidin (Sigma) for 5 days, then parasites were returned to drug-free media and media changed every 2-3 days until parasite population re-established. Genomic DNA was extracted using the PureLink genomic DNA mini kit (Invitrogen) and diluted to 10 ng/µl.

### Parasite growth assays

To test growth rates before RAP treatment, transgenic lines and parental parasites were synchronised at early rings, seeded at 200 µl in 96-well plates at 0.1% parasitaemia, 2% haematocrit and incubated for ∼72 h until the middle of the following cycle. 20 µl was taken for quantification by flow cytometry, then the media was changed, and parasites incubated for a further 24 h before quantification at the start of cycle 2 (after ∼96 h).

To test the phenotypes of RAP-treated parasites, ring stage parasites (4 h post-invasion) were synchronised with sorbitol and 0.05% DMSO or RAP (Sigma, 100 nM in DMSO, except PfELC-cKO: 200 nM) was added for 16 h. Cultures were washed twice in CCM and 200 µl dispensed in triplicate in 96 well plates at 1% parasitaemia, 0.3% haematocrit, with heparin-treated WT parasites (1:25, Pfizer) as a control. In each of the following three cycles, 100 µl was taken for flow cytometry and the remainder was diluted to 1% parasitaemia and incubated further. At the end of the first cycle, (∼40 h post-treatment), samples were taken for genotyping and Western blot analysis. For comparison of phenotypes under suspension and static conditions, after incubation with DMSO or RAP cultures were plated out in triplicate in 48 well plates in 150 µl at 5% haematocrit, 1% parasitaemia. This small volume supports consistent suspension of the culture. Identical plates were prepared and one incubated in a static incubator, the other incubated in a humidified box on a platform shaking at 185 rpm. 72 h post-treatment, 8 µl of culture was transferred to a 96 well plate and quantified by flow cytometry.

For flow cytometry analysis, 100 µl of parasites at 0.3% haematocrit was added to a 96 well plate and stained with SYBR Green I (Sigma, 1:5000) in 100 µl (15 minutes, room temperature) then washed three times in PBS and resuspended in 100-150 µl PBS for quantification. Flow cytometry was performed using a LSRFortessa cytometer (BD Biosciences) with high throughput sampler, with capture of 100,000 events per well. Samples were gated for RBCs, single cells then SYBR+ cells (Fig SX) and data were analysed using FlowJo (BD Biosciences), with each sample normalised to DMSO-treated control in each cycle.

### Microscopy analysis of parasites

For live fluorescence microscopy, late schizonts of prMA-GFP were stained with DRAQ5 DNA stain (Thermo Fisher, 5 µM, 30 minutes). The culture was resuspended in PBS for imaging, at a final haematocrit of 0.5%.200 µl was added to a well of an 8-well imaging slide (Ibidi, untreated) and allowed to settle. Images were acquired with an OrcaFlash 4.0 CMOS camera using a Nikon Ti Microscope (Nikon Plan Apo 60x or 100x 1.4-N.A. oil immersion objectives). Subsequent image manipulations were carried out in Fiji (Schindelin *et al*, 2012, 2015).

For video microscopy, DMSO and RAP-treated parasites were incubated for ∼40 h, then schizonts were isolated on gradients of 70% Percoll (Radfar *et al*, 2009) then washed in 10 ml CCM and the pellet size estimated. Isolated schizonts (>90% parasitaemia) were resuspended to 1% haematocrit in CCM and treated with egress inhibitor ML10 (Baker *et al*, 2017) at 1 µM for 3-5 h to synchronise at very late schizonts. Once mature, the culture was resuspended in CCM with fresh RBCs at 0.2% haematocrit, ∼50% parasitaemia. One at a time, samples were washed four times in warm CCM then resuspended in PBS for imaging and 200 µl added to an 8-well imaging slide and allowed to settle at 37°C. Samples were imaged using a Nikon Ti Microscope, 60x objective, enclosed within a heated incubation chamber, using a field of view with moderate density of cells. Egress begins 10-15 minutes after the final wash, and a 10-minute brightfield video (3 fps) was captured once a small fraction of schizonts had already egressed, to capture the most events. Immediately before and after the brightfield video, one frame of GFP fluorescence was also captured to assign schizonts as GFP+ or GFP-.

To quantify individual invasion attempts, each individual RBC that underwent echinocytosis was processed using Fiji. Invasion attempts were assigned to an event type based on successful invasion (merozoite clearly internalised, Type A), failed completion of internalisation (merozoite fully internalised, but ejected before the end of the video, Type B), failed initiation of internalisation (merozoite deforms RBC but is not internalised, Type C) or failed deformation (merozoite stably attached but does not deform or enter RBC, Type D). Deformation scores were assessed using the scheme of (Weiss *et al*, 2015), using a score of 0 to indicate no deformation and judging only the final deformation before invasion (if present).

For successful invasion and Type B failures, the phases of invasion were timed using an adjusted scheme from (Gilson & Crabb, 2009), starting from the first deformation (or stable attachment if no deformation present) and including end of deformation, start and end of internalisation and the start and maximal extent of echinocytosis. Events were excluded from analysis if any of the phases were obscured by other cells, the edge of the frame or the start or end of the video. RAP-treated parasite events were excluded if GFP-, except for PfELC-cKO which did not express GFP after truncation.

### Protein and immunochemistry techniques

For Western blot analysis, 5-10 ml of schizonts at <5% parasitaemia were lysed using 0.1% saponin/PBS (Sigma) for 10 min (room temperature), washed twice in PBS and lysed using RIPA buffer (Thermo Fisher). PfMyoB-cKO parasites were treated with E64 (Sigma, E3132) at 10 µM for 4-6 h to obtain as mature as possible schizonts before lysis. Parasite lysates were spun to isolate the soluble fraction (15000xg, 10’) and the supernatant was boiled with SDS for 5 min. When indicated, protein concentration was normalised between samples using the Pierce BCA protein assay kit (Thermo Fisher) before addition of SDS buffer. Samples were separated by SDS-PAGE using 4-12% Bis-Tris gels in MES buffer (Thermo Fisher) then stained with Coomassie or dry-transferred to a nitrocellulose membrane (iBlot2, Thermo Fisher) for Western blot. Blots were blocked and stained in 3% skimmed milk powder/PBST (0.1% Tween-20 in PBS), using anti-FLAG (1:2000, F1804, Sigma), anti-GFP (1:500, 7.1/13.1, Roche), anti-3xHA (1:2000, 12CA5, Roche or 1:4000, C29F4, Cell Signaling) anti-PfALD (1:1000, (Baum *et al*, 2006)), or anti-PfADF1 (1:2000, (Wong *et al*, 2011)) and HRP-coupled goat anti-mouse or -rabbit secondary antibody (1:5000, STAR120P/STAR121P, Bio-Rad). sBlots were washed in PBST and detected using ECL reagent (Amersham) and exposure to X-ray film or ChemiDoc imaging system (Bio-Rad).

