## Supplementary Figures and Tables for "Actomyosin forces and the energetics of red blood cell invasion by the malaria parasite *Plasmodium falciparum*"

#### Supplementary Information

**Includeed in this file:**

Supplementary Figures S1-S3

Supplementary Video S4-S8 legends

Supplementary Tables S1-S2

#### Supplementary Figures

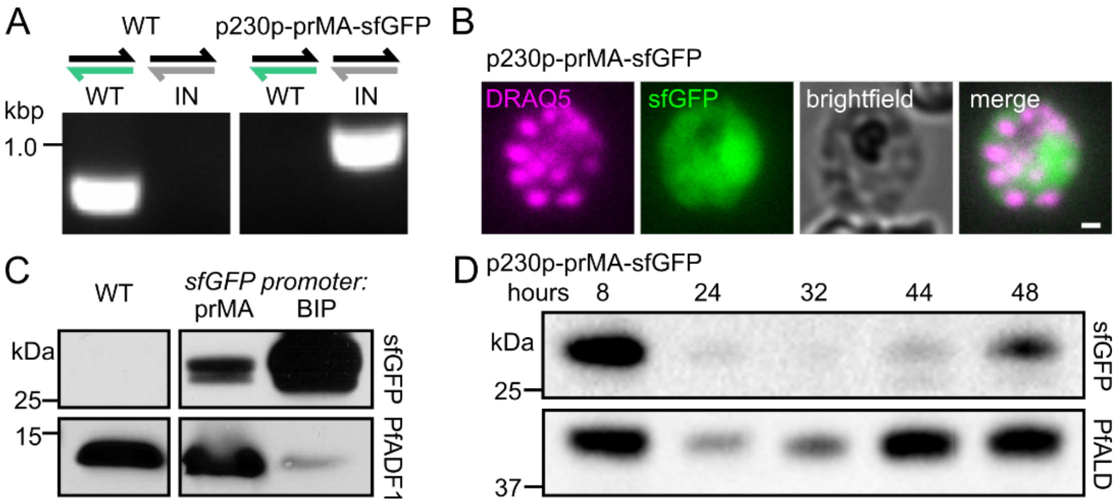


Figure S1: Validation of *Pfmyoa* promoter system at the *p230p* locus

**A** Exchanging the constitutive *BIP* promoter for the *Pfmyoa* promoter produced p230p-prMA-sfGFP parasites. Genotyping PCR showed a complete loss of the WT *p230p* locus (green half arrow) and gain of the integrated locus (IN, grey half arrow). **B** Live fluorescence microscopy of p230p-prMA-sfGFP schizont showing DNA (DRAQ5, magenta) and cytosolic sfGFP expression. **C** Western blot of p230p-prMA-sfGFP parasites confirming sfGFP expression, although at a lower level than constitutive p230p-BIP-sfGFP. **D** Western blot of p230p-prMA-sfGFP parasite lysate, normalised by total protein level, at different stages of the asexual life cycle. Strong expression of sfGFP was present in late schizonts and early rings only, as expected for the *Pfmyoa* promoter, while control protein *P. falciparum* aldolase (PfALD) levels dip slightly in trophozoite stages.

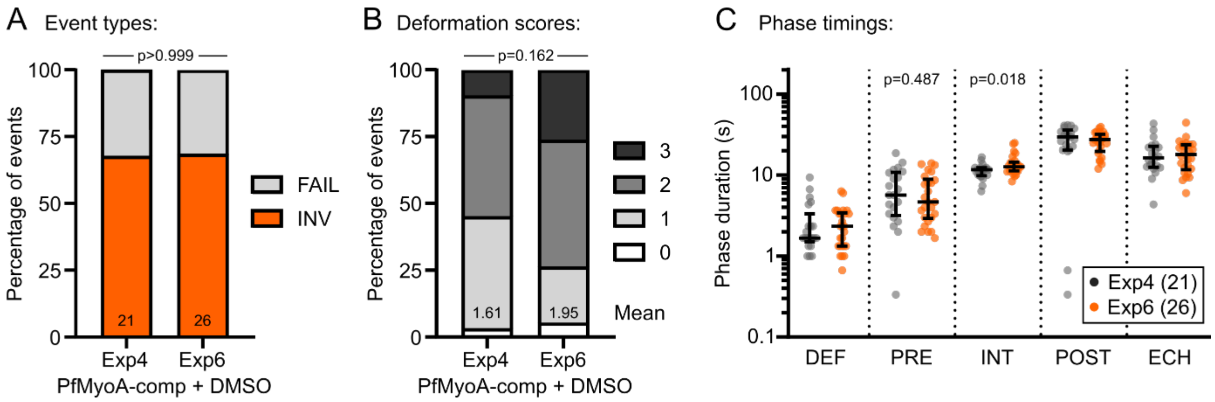


Figure S2: Video microscopy of PfMyoA-comp, broken down by biological repeat to assess consistency

**A** To assess the consistency of the two biological repeats, each two technical repeats, performed for each line, since the statistical tests assume that all events are independent, PfMyoA-comp + DMSO was chosen as an example where both repeats (Exp4 and Exp6) had similar numbers of events. The proportion of successful invasion events in both experiments was the same ($p>0.999$, Fisher's exact test, comparing pooled failures to Type A). **B** Comparing the deformation scores across the two experiments showed some differences, but not significant ($p=0.162$, chi square test). **C** Comparing the distributions of each phase duration across the two experiments showed only small differences. The duration of internalisation was significantly higher in Exp6, but the difference was small (11.7 s to 12.7 s). Bars show median and interquartile range. Significance assessed for each phase by Mann-Whitney test, shown when $p<0.5$.

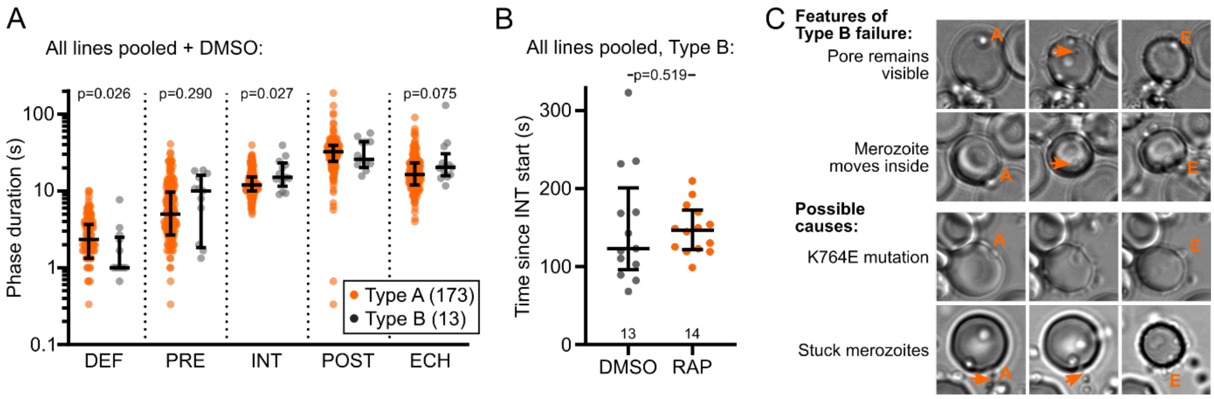


#### Figure S3: Type B events show slower internalisation

**A** From all lines after DMSO treatment, comparing Type A events to Type B events shows a significantly shorter deformation period and slower internalisation, with a trend to a longer pause before internalisation, consistent with either a merozoite producing less force or an RBC providing more resistance to invasion. Bars show median and interquartile range. Significance assessed for each phase by Mann-Whitney test, shown when p<0.5. **B** Comparing Type B failures from all DMSO- or RAP-treated lines shows no difference in the delay from start of internalisation to merozoite ejection. Bars show median and interquartile range. Significance assessed by Mann-Whitney test. **C** Example images of Type B failure where the invasion pore remained visible, hence not resealed (A= attachment, E= ejection); the incompletely-internalised merozoite continuing to move in a swirling pattern within the invagination; an ejection from PfMyoA-K764E after RAP treatment, where Type B failure was more common; and an event where resealing after internalisation was prevented by tethering to a sister merozoite.

### Supplementary Videos

#### Video S4: PfMyoA-cKO merozoites attach but do not deform or internalise

An example of a RAP-treated PfMyoA-cKO merozoite being stably attached (A) but neither deforming nor being internalised before echinocytosis (E), as expected from the absence of the PfMyoA motor. Time indicated in seconds, scale bar 2 µm.

#### Video S5: PfELC-cKO merozoites deform but do not internalise

An example of a RAP-treated PfELC-cKO merozoite showing attachment (A) and deformation (D) but no internalisation before echinocytosis (E), indicative of a weakened, but present, PfMyoA motor. Time indicated in seconds, scale bar 2 µm.

#### Video S6: PfMyoA-K764E merozoites are more often ejected after internalisation

An example of a RAP-treated PfMyoA-K764E merozoite showing apparently normal attachment (A), deformation (D) and internalisation (I), before the merozoite was subsequently ejected (arrowhead), after echinocytosis (E) was completed. This suggests a weakened PfMyoA motor, but the merozoite appeared to remain motile while emerging from the RBC. Time indicated in seconds, scale bar 2 µm.

#### Video S7: PfMyoB-cKO merozoites take longer to initiate internalisation

An example of a RAP-treated PfMyoB-cKO merozoite showing apparently normal attachment (A), deformation (D) and internalisation (I), before echinocytosis (E). On average the loss of PfMyoB delayed the start of internalisation. Time indicated in seconds, scale bar 2 µm.

#### Video S8: *P. falciparum* merozoites exhibit swirling motility before and during invasion

An example of a RAP-treated PfMyoB-cKO merozoite (though WT merozoites behaved similarly), demonstrating a swirling motion (arrowhead) after attachment (A), before internalisation (I) and echinocytosis (E). This matches observations of *P. knowlesi* merozoites, which glide in a corkscrew motion on a substrate (Yahata *et al*, 2020). Time indicated in seconds, scale bar 2 µm.

### Supplementary Tables

#### Table S1: Oligonucleotides used in this study for genotyping PCR and sequencing

| # | Locus | Part | Sequence | Source |
|---|---|---|---|---|
| 1 | *p230p* | WT[a]/IN[b]-F | ACCATCAACATTATCGTCAG | Ashdown *et al, in press* |
| 2 | *p230p* | WT-R | TCTTCATCAGCCTGGTAAC | Ashdown *et al, in press* |
| 3 | *p230p* | IN-R | CATTTACACATAAATGTCACAC | Ashdown *et al, in press* |
| 4 | PfMyoA-K764E | IN-F | GTTCATTGATTGGATCTCAGTTC | This study |
| 5 | PfMyoA-K764E | IN-R | GGATACGAGCCAGCATAGTC | This study |
| 6 | PfMyoA-cKO | IN/EX[c]-F | GGTCGTTTCATGCAGTTGGT | (Robert-Paganin *et al*, 2019) |
| 7 | PfMyoA-cKO | IN-R | GCCAGCCACGATAGCCGCGCTGCCTCGTCCTGCAGT TCATTCAGGGCACCGGACAGGTCG | (Robert-Paganin *et al*, 2019) |
| 8 | PfMyoA-cKO | EX-R | ACCTTCACCCTCTCCACTGAC | (Robert-Paganin *et al*, 2019) |
| 9 | PfMyoB-cKO | WT/IN/EX-F | ATGGGATCGAAAAGGGTGGT | This study |
| 10 | PfMyoB-cKO | WT-R | TCCATGATCACTCGTCCTCAC | This study |
| 11 | PfMyoB-cKO | IN-R | AAGTCTCCACAATTGATAAAGAG | This study |
| 12 | PfMyoB-cKO | EX-R | TTATTTGTACAGTTCATCCATACC | This study |
| 13 | pDC2-p230p-BSD | Seq[d] (*bsd*) | TTTTTGTAATTTCTGTGTTTATG | This study |
| 14 | pDC2-generic | Seq (gRNA) | AAGCACCGACTCGGTGCCAC | Marcus Lee |
| 15 | p230p-prMA-sfGFP | Seq (*sfgfp*) | TGAACCATACGGGTTGTTG | This study |
| 16 | p230p-prMA-MyoA | Seq (*myoa*) | CTCCTGGAGCCAAGCAC | This study |

[a]WT = wild type locus, [b]IN = integrated locus, [c]EX = excised locus, [d]Seq = sequencing

#### Table S2: Oligonucleotides used in this study for cloning PCR and guide RNAs

| # | Construct | Part | Sequence | Source |
|---|---|---|---|---|
| 17 | pDC2-p230p-BSD | BSD-F | ATATCCATGGCCAAGCCTTTGTCTCAAGAAGAATCC | This study |
| 18 | pDC2-p230p-BSD | BSD-R | ATATCCGCGGTTAGCCCTCCCACACATAAC | This study |
| 19 | p230p-prMA-sfGFP | prMA-F | ATATCCGCGGAGCATGCACAATTAAGAAGACG | This study |
| 20 | p230p-prMA-sfGFP | prMA-R | ATATGCTAGCTTTTTTTTTTTTTTTTATAAATATGAA AAG | This study |
| 21 | p230p-prMA-PfMyoA | MyoA-F | CTTTTCATATTTATAAAAAAAAAAAAAAAAGCTAGCA TGGCCGTTACCAACG | This study |
| 22 | p230p-prMA-PfMyoA | MyoA-R | GATATTTACTTATTTATTTTATTCTGCATATTTAAA AATCCTGCAGTTAGACCGCATAATCCGG | This study |
| 23 | p230p-prMA-K764E | SDM[a]-F | GCAAGAGGGTGCTGAAATTTTAACAAAAATAC | This study |
| 24 | p230p-prMA-K764E | SDM-R | ATTTTTGTTAAAATTTCAGCACCCTCTTGC | This study |
| 25 | p230p-prMA-S19A | SDM-F | GTGAGGAGAGTAGCTAACGTGGAGG | This study |
| 26 | p230p-prMA-S19A | SDM-R | CAAAAGCCTCCACGTTAGCTACTCTCCTC | This study |
| 27 | p230p-prMA-DN | MyoA-F | CTTTTCATATTTATAAAAAAAAAAAAAAAAGCTAGCA TGAACGTGGAGGCTTTTGATAAATC | This study |
| 28 | pUC-PfMyoB-CK | HR[b]-F | CAGTCACGACGTTGTAAAACGACGGCCAGTGAATTC TTAGAGAATTTCATAGGTATTTTGG | This study |
| 29 | pUC-PfMyoB-CK | HR-R | TGCGTAATCCGGTACATCATATGGGTACATTTCATG CTCTTTTATATATTTGTACTTAC | This study |
| 30 | pUC-PfMyoB-CK | SF[c]-F | ATGTACCCATATGATGTACCG | This study |

| | | | | |
|---|---|---|---|---|
| 31 | pUC-PfMyoB-CK | SF-R | TTATTTGTACAGTTCATCCATACC | This study |
| 32 | pUC-PfMyoB-CK | 3UTR[d]-F | ACGCATGGTATGGATGAACTGTACAAATAACTCGAG AAATCGGGAAAATAAAATGG | This study |
| 33 | pUC-PfMyoB-CK | 3UTR-R | TATGACCATGATTACGCCAAGCTTGCATGCCTGCAG GTCGTCCCAATCATTTTTC | This study |
| 34 | pDC2-p230p-hDHFR | gRNA[e]-F | TATTAGGCTGATGAAGACATCGGG | Ashdown *et al, in press* |
| 35 | pDC2-p230p-hDHFR | gRNA-R | AAACCCCGATGTCTTCATCAGCCT | Ashdown *et al, in press* |
| 36 | pDC2-pfmyob-hDHFR-1 | gRNA-F | TATTGTATAAGATAGGAAAAAGCA | This study |
| 37 | pDC2-pfmyob-hDHFR-1 | gRNA-R | AAACTGCTTTTTCCTATCTTATAC | This study |
| 38 | pDC2-pfmyob-hDHFR-2 | gRNA-F | TATTGTAGAAGCATTTGACAAAAG | This study |
| 39 | pDC2-pfmyob-hDHFR-2 | gRNA-R | AAACCTTTTGTCAAATGCTTCTAC | This study |

[a]SDM = site-directed mutagenesis, [b]HR = homology region, [c]SF = synthetic fragment, [d]3UTR = 3' un-translated region, [e]gRNA = guide RNA.
